# Deep genome sequencing and variation analysis of 13 inbred mouse strains defines candidate phenotypic alleles, private variation, and homozygous truncating mutations

**DOI:** 10.1101/039131

**Authors:** Anthony G. Doran, Kim Wong, Jonathan Flint, David J. Adams, Kent W. Hunter, Thomas M. Keane

## Abstract

**Background:** The Mouse Genomes Project is an ongoing collaborative effort to sequence the genomes of the common laboratory mouse strains. In 2011, the initial analysis of sequence variation across 17 strains found 56.7M unique SNPs and 8.8M indels. We carry out deep sequencing of 13 additional inbred strains (BUB/BnJ, C57BL/10J, C57BR/cdJ, C58/J, DBA/1J, I/LnJ, KK/HiJ, MOLF/EiJ, NZB/B1NJ, NZW/LacJ, RF/J, SEA/GnJ and ST/bJ), cataloging molecular variation within and across the strains. These strains include important models for immune response, leukemia, age-related hearing loss and rheumatoid arthritis. We now have several examples of fully sequenced closely related strains that are divergent for several disease phenotypes.

**Results:** Approximately, 27.4M unique SNPs and 5M indels are identified across these strains compared to the C57BL/6J reference genome (GRCm38). The amount of variation found in the inbred laboratory mouse genome has increased to 71M SNPs and 12M indels. We investigate the genetic basis of highly penetrant cancer susceptibility in RF/J finding private novel missense mutations in DNA damage repair and highly cancer associated genes. We use two highly related strains (DBA/1J and DBA/2J) to investigate the genetic basis of collagen induced arthritis susceptibility.

**Conclusion:** This paper significantly expands the catalog of fully sequenced laboratory mouse strains and now contains several examples of highly genetically similar strains with divergent phenotypes. We show how studying private missense mutations can lead to insights into the genetic mechanism for a highly penetrant phenotype.

## Background

Genetic mutations (such as SNPs, indels and structural variants) are commonly involved in the dysregulation of normal gene function and frequently the cause of human diseases. Laboratory mouse strains are one of the primary model organisms used to study human disease, and have excelled as a model system in which to identify and characterise the role that genomic variation plays in susceptibility and pathogenesis [1,2]. A multitude of inbred laboratory strains are now commonly employed in the examination of a wide range of human diseases [3].

The first draft version of the mouse reference genome was produced based on a whole genome shotgun sequencing strategy of a female C57BL/6J strain mouse [4] followed several years later by the finished genome sequence [5]. The mouse reference genome enabled a whole set of new applications and technologies such as many new genetic screens [6], the creation of null alleles for the majority of genes [7], and accelerated the discovery of genetic diversity in other laboratory mouse strains [8]. The Mouse Genomes Project (MGP) was initiated with the primary goals to sequence the genomes of the common laboratory mouse strains, catalogue molecular variation and produce full chromosome sequences of each strain [9]. The initial phase of the MGP focused on sequencing and variation analysis of a representative subset of the most commonly used inbred mouse strains. This culminated in the identification of 57.7M SNPs, 8.8M indels and 0.71M structural variants (SV) across the genomes of 17 of the most commonly used strains [9].

Here, we present a significant expansion of the MGP variation catalogue by deep sequencing (30–60x) and variation analysis of 13 strains: BUB/BnJ, C57BL/10J, C57BR/cdJ, C58/J, DBA/1J, I/LnJ, KK/HiJ, MOLF/EiJ, NZB/B1NJ, NZW/LacJ, RF/J, SEA/GnJ and ST/bJ. These represent a diverse set of phenotypes and genotypes of inbred mouse strains that are commonly used to study rheumatoid arthritis (DBA/1J), diabetes (KK/HiJ), immunity (C57BL/10J, NZB/B1NJ, NZW/LacJ), cancer (C58/J, RF/J), development (I/LnJ, SEA/GnJ) and age-related diseases (BUB/BnJ, C57BR/cdJ, MOLF/EiJ, ST/bJ). The catalogue of variants derived from these strains has been incorporated into the current MGP variation catalogue to determine private and shared variation among the 13 strains, and refine private variation identification across all strains in the catalogue.

We also demonstrate how variants in these 13 strains can be utilised to investigate the genetic basis of disease. In the first analysis, predicted damaging SNPs private to the RF/J strain were used to identify candidate genes and biological pathways involved in tumorigenesis and disease progression. The RF/J strain is an important model in cancer studies due to a naturally high predisposition to several cancers including acute myeloid leukemia and thymic lymphoma [10,11]. The second analysis involved the identification of candidate genes for collagen induced arthritis (CIA) susceptibility, a model of human rheumatoid arthritis (RA), by comparison of SNPs between two genetically related strains with divergent phenotypes. Candidate genes identified in the analysis of these two strains, DBA/1J (susceptible) and DBA/2J (resistant), were then further investigated to identify potential causal pathways and biological processes controlling pathogenesis of this disease.

## Results and Discussion

### A phenotypically diverse set of laboratory mouse strains

A total of 13 genetically and phenotypically diverse inbred laboratory mouse strains were chosen for analysis in the current study (Fig. 1). These strains were selected from across the inbred mouse subspecies groups and represent a large number of model systems used to study human disease. This data set includes strains that are genetically similar to the current MGP strains, such as DBA/1J and C57BL/10J, as well as strains that are genetically divergent, such as the wild-derived Japanese strain MOLF/EiJ (Fig. 1).

**Figure 1.**
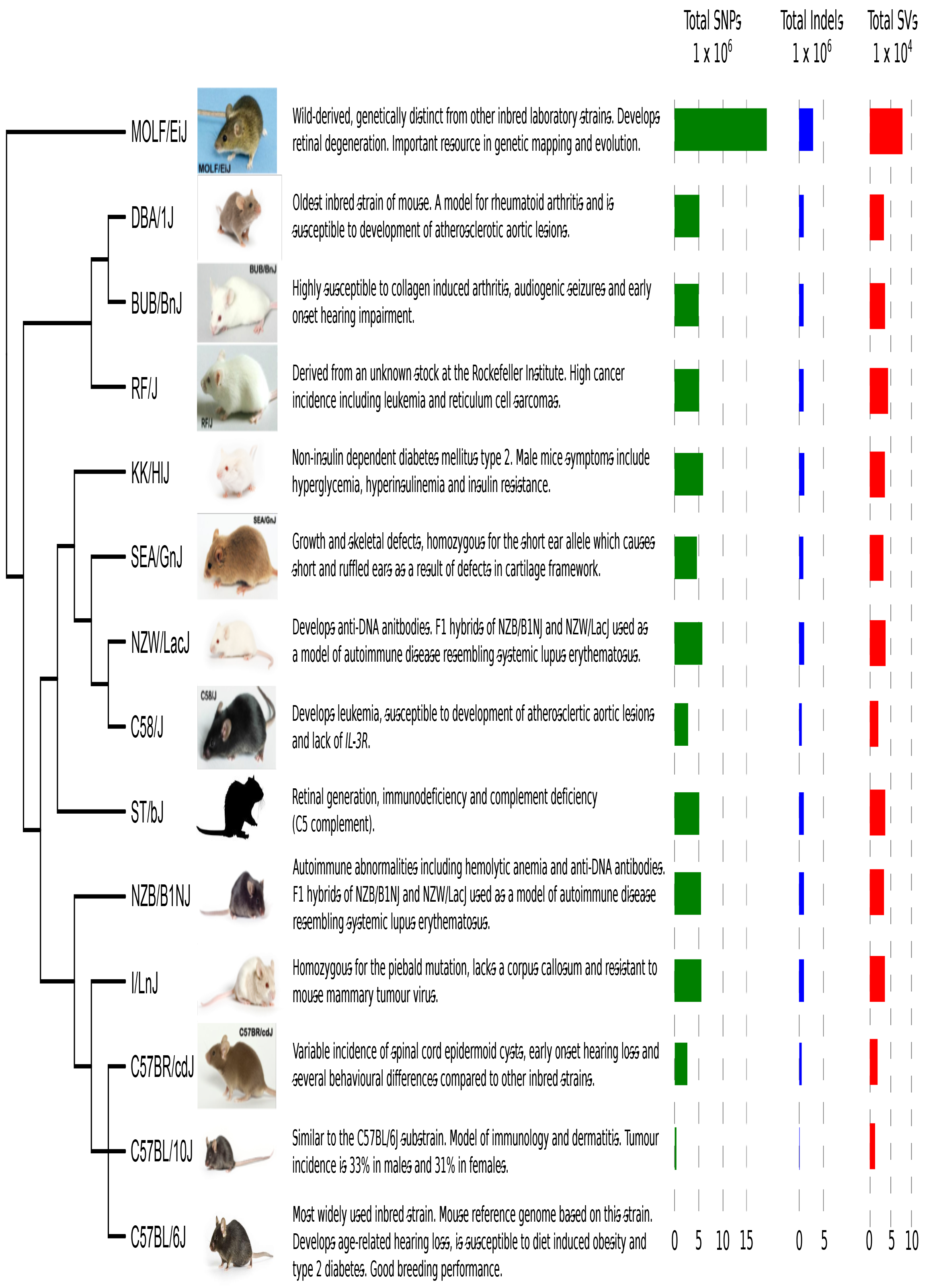
**Phenotypes and variant statistics for each of the 13 strains.** The strains sequenced in this project with a short description and notable phenotypes, followed by the total number of SNPs, indels and large deletions. Individual strain images provided by The Jackson Laboratory (ME, USA). The phylogenetic tree is a genome-wide summary built using all of the HapMap genotypes for each strain [22] and does not reflect local haplotype structure.

With this expansion, there are now several examples of fully sequenced strains which are closely related to other strains in the catalogue but are divergent for disease phenotypes. For example, C57BR/cdJ mice are closely related to the mouse reference genome strain, C57BL/6, but display different patterns of learning behaviour [12]. Following avoidance training, C57BL/6 mice exhibit poor conditioned avoidance performance, whereas C57BR/cdJ perform much better in avoidance tests [13]. Another example of genetically similar but phenotypically divergent strains include DBA/1J and DBA/2J which are susceptible and resistant, respectively, to collagen induced arthritis. DBA/1J mice are commonly used as a model for rheumatoid arthritis and exhibit several similarities to the human disease including synovitis and degradation of the bone and cartilage tissue [14].

Among this expansion of the variation catalogue are several strains with similar phenotypes such as the NZB/B1NJ and NZW/LacJ strains. These strains are widely used as models for autoimmune abnormalities. For example, both strains develop anti-DNA antibodies. Also, F1 hybrids of these two strains develop an autoimmune disease resembling human systemic lupus erythematosus [15].

Another interesting addition to the variation catalogue examined here is the RF/J strain [10,11]. This strain is an important model for cancer research due to a naturally high incidence rate of acute myeloid leukemia and tumour development. This strain also exhibits a high spontaneous rate of glomerular hyalinisation and glomerulosclerosis [16]. Each of the remaining strains also exhibit several notable phenotypes; BUB/BnJ are homozygous for the *Mass1^frings^* allele, which is responsible for early onset (3–4 weeks of age) hearing impairment [17]; a high proportion of I/LnJ mice display agenesis of the corpus callosum which is associated with developmental delays and impaired motor function in humans [18]; SEA/GnJ mice segregate for the short ear allele *(BMP5^se^),* which causes numerous defects in cartilage and bone tissue [19]; both the ST/bJ and MOLF/EiJ strains are homozygous for the retinal degeneration allele *Pde6b^rd1^* [20] [11853768]; and C58/J and KK/HiJ mice are homozygous for the age related hearing loss mutation, *Cdh23^ahl^* [21].

### Identification of SNPs and indels

The 13 mouse strains were sequenced to a mapped depth of 30–60x on the Illumina HiSeq platform at a read length of 101 bp paired-end (Table 1). In total, over 1. 8Tb of raw sequencing data was generated. All raw sequencing reads are available from the European Nucleotide Archive (ERP000927; see Table s1 in Additional file 1). Sequencing reads were aligned to the GRCm38 mouse reference genome using BWA-mem, followed by a local realignment around indels using GATK. The aligned sequence reads are also available from the Mouse Genomes Project (ftp://ftp-mouse.sanger.ac.uk).

**Table 1.** **Sequencing, alignment and variant statistics.** Total sequencing in Gigabases (Gb), mapped coverage (based on 3Gb genome) and the total and private number of SNPs, indels and large deletions is shown.

| Strain | Raw bases (Gb) | Mapped Coverage | SNPs | private | Indels | private | SVs | Private |
| --- | --- | --- | --- | --- | --- | --- | --- | --- |
| BUB/BnJ | 135.42 | 38.92 | 4982043 | 54418 | 929617 | 19187 | 37034 | 5375 |
| C57BL/10J | 108.43 | 29.85 | 349702 | 2733 | 83930 | 1893 | 12939 | 9904 |
| C57BR/cdJ | 140.94 | 39.76 | 2601138 | 9683 | 485803 | 6307 | 18927 | 3462 |
| C58/J | 153.06 | 43.76 | 2834841 | 27805 | 547151 | 11726 | 20666 | 3777 |
| DBA/1J | 139.79 | 38.55 | 5129920 | 9001 | 955621 | 9684 | 34239 | 4480 |
| I/LnJ | 124.97 | 34.11 | 5547267 | 78583 | 1013686 | 24628 | 36612 | 6342 |
| KK/HiJ | 151.60 | 44.42 | 5893784 | 94110 | 1078555 | 31084 | 36588 | 3725 |
| MOLF/EiJ | 112.55 | 30.68 | 19027669 | 1818029 | 2892638 | 367208 | 79239 | 17394 |
| NZB/B1NJ | 133.24 | 36.15 | 5473276 | 54189 | 1014782 | 20670 | 34406 | 3164 |
| NZW/LacJ | 160.46 | 49.06 | 5736044 | 52902 | 1051170 | 21397 | 38207 | 3308 |
| RF/J | 147.33 | 43.18 | 5090115 | 32390 | 937091 | 13440 | 44205 | 11685 |
| SEA/GnJ | 134.23 | 38.67 | 4603720 | 30263 | 857599 | 12729 | 33033 | 3686 |
| ST/bJ | 223.17 | 63.74 | 5107777 | 70064 | 978238 | 24325 | 37350 | 4474 |

A total of 27.39M high confidence homozygous SNPs were identified across the genomes of 13 inbred laboratory mouse strains (Fig. 1). This catalogue of variants, when compared to the MGP variation catalogue of 23 strains, contains 2.78M (10.15%) novel variant sites, of which 2.33M (8.51%) are private to one of the 13 strains in this study when compared to the entire variation catalogue (Fig. 2; Additional file 2). The wild-derived MOLF/EiJ strain contained the most individual variation both in terms of total (19.03M) and private SNPs (1.82M). Conversely, the C57BL/10J strain, which is genetically much more similar to the reference genome, displayed the least amount of total and private variation (349,702 and 2,733). The total number of SNPs and short indels shared between each pair of strains contained in the MGP variation catalogue is given in Additional file 3.

**Figure 2.**
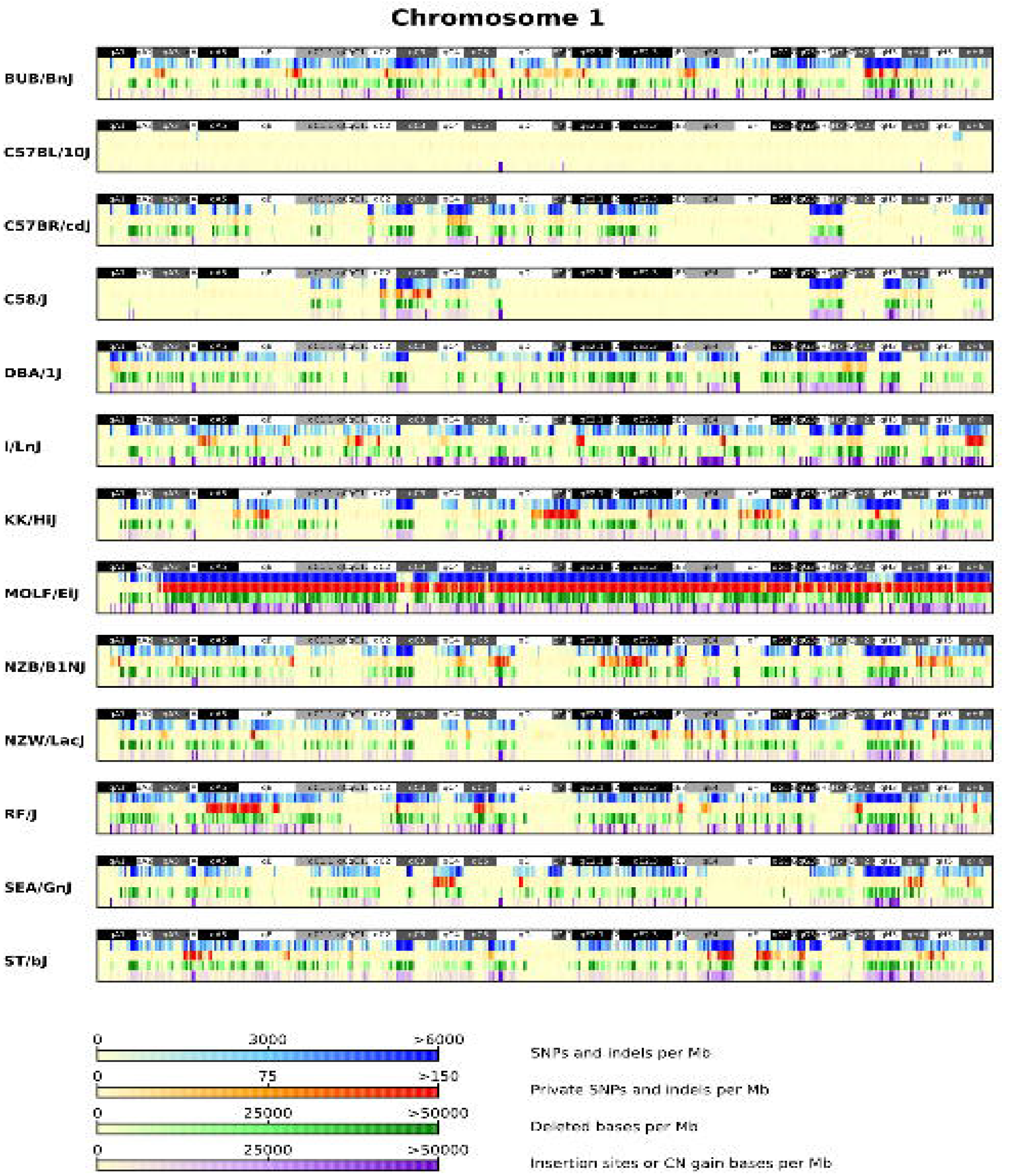
**SNP, indel and SV densities for chromosome 1 of all strains.** SNP, indel and SV (insertions and deletions) densities (per MB) for all variants identified on chromosome 1 for each of the 13 strains.

In total 4.81M high quality homozygous indels were identified, 0.64M of which are novel indels sites, and 0.56M of these are private to one of the 13 strains when compared to the all other strains included in the MGP variation catalogue (Fig. 2; Additional file 2). The 2.91M deletions identified range in size from 1 bp to 46 bp, and the 2.5M insertions range in size from 1 bp to 29 bp. Similar to the number of SNPs identified, MOLF/EiJ contained the largest number of short indels relative to the reference genome (2.89M total and 367,208 private). Additional comparisons of the total number of indels shared between each pair of strains is contained in (Additional file 3).

To ensure the accuracy of our SNP calling pipeline, genotypes from the mouse HapMap [22] for approximately 138,000 locations were obtained and compared to the corresponding genotypes in each of the 13 strains involved in this study. After removing missing genotypes and mapping genotype locations from NCBIm37 to GRCm38 of the reference genome, between 115,572 (MOLF/EiJ) and 129,825 (C58/J) genotypes remained for comparison with each strain. Sensitivity of our calling pipeline was determined by assessing the concordance rate for each of the individual strains. Greater than 98% of genotypes were concordant for each of the individual strains (Additional file 4). In fact, a total of 1.64M genotypes were compared, in which 1.62M (98.68%) were concordant. All of the variants in this study were incorporated into the MGP v5 variation catalogue and are available from the project website (ftp://ftp-mouse.sanger.ac.uk and the European Variation Archive (PRJEB11471). The MGP v5 catalogue comprises 36 inbred laboratory strains with 70.72M SNP sites, 39.23M of which are private to an individual strain, and 11.29M short indels, 6M of which are private.

### Functional consequences of SNPs and indels

The Variant Effect Predictor (VEP) tool [23] was used to assign functional annotations to the SNP and indel variants across the 13 strains (Table 2 and Additional file 5). The majority of SNPs are found in intergenic (46.58–52.28% across strains; 50.57% of all SNPs identified) and intronic (15.74–18.58%; 18.08%) regions. Additionally, a large amount of variation was identified up and downstream of protein coding genes (3.91–4.47%; 3.93% and 3.79–4.43%; 4.05%, respectively), and within the 3’ (0.20–0.24%) and 5’ (0.02–0.04%) untranslated regions (UTR).

**Table 2.** Predicted consequences of variants.

|  | SNPs |  |  | Indels |  | Exonic SVs |  |
| --- | --- | --- | --- | --- | --- | --- | --- |
| Strain | Synonymous | Non-synonymous | Stop gain/loss | Frameshift variant | Inframe variant | Deletions | Insertions |
| BUB/BnJ | 9751 | 6050 | 44 | 41 | 169 | 1247 | 720 |
| C57BL/10J | 581 | 298 | 0 | 4 | 8 | 57 | 102 |
| C57BR/cdJ | 4400 | 2948 | 18 | 26 | 52 | 288 | 227 |
| C58/J | 5238 | 3362 | 23 | 35 | 69 | 330 | 254 |
| DBA/1J | 9782 | 6198 | 39 | 53 | 149 | 419 | 389 |
| I/LnJ | 10389 | 6249 | 42 | 52 | 150 | 690 | 1671 |
| KK/HiJ | 10816 | 6197 | 50 | 56 | 140 | 599 | 462 |
| MOLF/EiJ | 33710 | 17311 | 108 | 172 | 439 | 897 | 1305 |
| NZB/B1NJ | 10403 | 6367 | 43 | 62 | 146 | 500 | 390 |
| NZW/LacJ | 10434 | 6326 | 46 | 61 | 146 | 673 | 425 |
| RF/J | 9891 | 6174 | 39 | 56 | 140 | 2033 | 862 |
| SEA/GnJ | 9309 | 6027 | 38 | 57 | 138 | 1284 | 614 |
| ST/bJ | 9021 | 5578 | 45 | 59 | 133 | 713 | 413 |

Although most SNPs were found in noncoding regions, a large amount of variation was located in protein coding regions (76,108 SNPs). Of particular note are SNPs that affect the protein coding sequence, such as splice variants and non-synonymous SNPs. A total of 27,416 (0.1%) missense SNPs were identified across the 13 strains. MOLF/EiJ contained the largest number of missense variants (17,311), however, only 2,288 of these were private to this strain. One particular novel missense mutation is a G→A transition in exon 4 of the *Pctp* gene identified in the NZW/LacJ and NZB/BlNJ strains. This mutation, at position 89,987,348 on chromosome 11, is also shared with the NZO/HlLtJ strain which suggests fixation of the mutation in progenitor mice prior to development of the NZ inbred mouse strains [24]. This mutation causes an arginine to histidine change at amino acid position 120, which leads to inactivation of the protein. Although the physiological implications of inactivating *Pctp* are not yet fully understood, inactivation has been linked to changes in control of hepatic cholesterol homeostasis [25] and impaired biliary lipid secretion [26].

We identified 178 SNPs that result in a stop gain or loss. MOLF/EiJ contained the most individual variation in coding regions including 23 stop losses and 85 stop gains, however only 1 stop loss and 18 stop gain variants were private to this strain.C57BL/10J was the only strain in which a stop gain or loss was not identified. Several mouse strains are known to carry homozygous mutations that lead to the introduction of a premature stop in the coding sequence. Among the list of stop gains identified in our study was a C→A mutation which causes a premature stop codon in the *Pde6b* gene on chromosome 5 at position 108,421,265. This mutation is found in several strains included in the MGP variation catalogue, including three from this study; BUB/BnJ, MOLF/EiJ and ST/bJ. This mutation, rs49995481, results in truncation of the predicted protein in which over half of the protein is missing and is known to cause early onset severe retinal degeneration [27].

To determine the impact of an amino acid substitution, SIFT (Sorting Intolerant from Tolerant) [28] and grantham matrix scores (GMS) [29] were estimated for all non-synonymous SNPs. SIFT scores ≤ 0.05 are usually considered to be indicative of a deleterious mutation. Although most non-synonymous SNPs are tolerated, 11,182 in 6,241 genes were predicted to have a deleterious effect. Similar to SIFT, GMS enables the identification of moderately radical (100<GMS≤150) and radical (GMS>150) non-synonymous substitutions. Only 3,337 (10.78%) of non-synonymous substitutions are considered radical using GMS, although 1,130 of these are also annotated as deleterious using SIFT.

Similar to SNPs, most indels are intergenic (47.76%), intronic (18.82%) or within 5kb upstream (4.26%) or downstream (4.27%) of a coding gene. Although, in general, indels found in intergenic or intronic regions are not known to have an effect on protein coding genes, several strains carry a 5bp deletion at the branching point of intron 7 of the *Il3ra* gene which leads to skipping of exon 8 [30]. This mutation, on chromosome 14 at position 14,350,904, is found in 9 MGP strains including 5 from this study; BUB/BnJ, C58/J, I/LnJ, NZB/BlNJ and RF/J. A small number of indels cause an in-frame deletion (368) or insertion (278), and only 308 indels cause a frameshift. As with missense SNPs, MOLF/EiJ contained the largest number of frameshift mutations (172), however, proportional to the total number of indels identified in this strain, this number was approximately the same as the other strains (0.0006%). Several strains are known to carry indels that affect the protein coding function of genes, such as a known TA deletion in the hemolytic complement gene. This deletion, at position 35,043,207 on chromosome 2, leads to the creation of a premature stop codon 4 bases after the deletion [31] and consequently macrophages from mice carrying this mutation fail to secrete complement 5 (C5) [32]. This mutation is carried by several strains in the MGP catalogue, including 6 strains from this study; I/LnJ, KK/HiJ, MOLF/EiJ, NZB/B1NJ, RF/J and ST/bJ. Mice deficient for complement 5 have previously demonstrated enhanced protection against development of severe glaucoma compared to mice of the same strain sufficient for C5 [33].

### Novel and private variation

To identify variants unique to an individual strain (private SNPs and indels), genotypes for all variant sites identified in each of the 13 strains were computationally derived for all strains in the MGP variation catalogue (36 strains). Private SNPs were then identified as high quality homozygous alternative sites, in which the same site must be a high quality homozygous reference allele in at least 30 other strains. A maximum of 5 strains were allowed a missing genotype or low quality call at the same site to account for the genetic diversity of wild-derived strains in the MGP variation catalogue. This resulted in 2.33M (8.52%) private SNPs across the 13 strains, although 1.82M of these private sites were identified in the wild-derived MOLF/EiJ strain (Table 1). This accounted for 9.56% of all MOLF/EiJ SNP sites, which was much higher than the proportion of private variation identified for any of the other 12 strains (0.18–1.6%) (Table 1). DBA/1J had the lowest proportion of private SNPs due to the genetic similarity of DBA/1J and DBA/2J [34]; 90.72% of DBA/1J SNPs are shared with DBA/2J.

Similar to the total variation per strain, most private SNPs are located in intergenic (45.67–57.68% per strain) and intronic regions (13.28–23.32%) while 7.18–10.3% per strain are within 5kb of a coding gene. In total, 7,237 private missense SNPs were identified in 4,615 genes across all 13 strains although 4,086 of these are estimated to have more than one consequence (Additional file 6). Most private missense SNPs are predicted to be tolerated by SIFT or have a GMS < 100. Using SIFT predictions, 2,146 SNPs affecting 1,768 genes were predicted to be deleterious. Similarly, using GMS predictions, only 893 SNPs affecting 818 genes were predicted to cause moderate changes and 461 SNPs affecting 446 genes were predicted to cause radical changes to the CDS. As expected, only a small number of private variants led to changes in stop codons (93 stop gains and 12 stop losses) or were predicted to affect splice acceptor or donor sites (46 and 53 SNPs, respectively) although many of these SNPs were predicted to have more than one consequence (51 stop gains, 8 stop losses, 43 splice acceptor and 43 splice donor sites).

Ninety-one genes are affected by a private stop gain. For example, a C→T transition at amino acid position 208 of the bone morphogenetic protein 5 *(BMP5)* gene causes a stop gain private to the SEA/GnJ strain. Bone morphogenetic proteins are signalling molecules involved in the development of bone and cartilage tissue [35]. This strain is known to be homozygous for this mutation (also known as the short ear allele). Mice homozygous for the short ear allele exhibit abnormal skeletal development leading to smaller and weaker bones, reduced shape and size of the external ear, a reduced number of ribs and alterations in size and shape of many small bone elements [36].

Nearly half of private indels (48.96%) are found in intergenic regions, 18.61% are intronic and 8.89% are within 5kbp of a coding gene while almost a quarter (23.27%)-of indels are predicted to have 2 or more consequences. Only 84 (0.015%) private frameshift mutations were identified, and 61 of these were identified in MOLF/EiJ. As expected, very little private variation was observed in the C57BL/10J strain compared to the C57BL/6J reference genome. These two strains were separated in 1921 from the same C57BL progenitor colony, and no other strains are known to have contributed to the parentage of either strain [3]. Eighty-three genes were identified that carry a private frameshift mutation in one of the 13 strains such as a “T” insertion resulting in a frameshift at amino acid position 431 of the phosphorylase kinase alpha 1 (*Phka1)* gene. This mutation, which is private to the I/LnJ strain, causes a premature stop codon in the downstream coding region resulting in a great reduction of *Phka1* transcript in skeletal muscle and partial reduction in heart and brain [37]. Additionally, this mutation has been implicated in myopathy and glycogen storage disease in both humans and mice [38].

### Large structural deletions and insertions

A total of 79,943 large (>100 bp) deletions and 143,854 insertions were identified across all of the strains (Table 1). Of the large deletions identified, 20,122 were novel and a large proportion (17,514) of these were private to one of the 13 strains when compared to the full variation catalogue of 36 strains. Structural variant insertions were also identified for all strains in the MGP variation catalogue, including the 13 strains from this study. A total of 143,854 insertions were identified across the 13 strains examined in this study. Approximately 48.34% of these were novel (69,542), 63,262 of which were private to one of the 13 strains when compared to the MGP variation catalogue.

To assess the accuracy of our calling pipeline, PCR validation deletions and insertions were obtained for 7 MGP strains [39]. Structural variants were identified in these 7 strains using the same approach as for the 13 strains in this study. Sensitivity estimates were estimated for each of the 7 strains (Table S2 of Additional file 1). All sensitivity estimates for each strain were between 92.05–94.12% for deletions and 81.94-94.25% for insertions, which are comparable to estimates previously published for MGP strains [40,41].

Although, none of the 13 strains included in this study were among the 7 strains,151 PCR validated deletions and 84 insertions were available for the DBA/2J strain. The 151 PCR validated deletions identified in the DBA/2J strain [39] were compared with the deletion call set identified in DBA/1J. Although these are two separate strains, they are genetically identical at many loci throughout the genome. Most of the 151 deletions overlapped with a deletion identified in DBA/1J from our analysis (131). This meant that 20 deletions PCR validated in DBA/2J were not identified in the DBA/1J strain. Upon visual inspection of these deletions using the alignment data and the integrative genomics viewer [42], eight were identified as real differences between the strains. When true structural variant differences between the strains are removed from the sensitivity analysis, the sensitivity increases to 91.61% (131/143) which is similar to the proportion of SNP sites that are identical between the strains (90.72%).

Sensitivity estimates were also calculated for SV insertions identified in the DBA/1J strain using PCR validated calls identified in the DBA/2J strain. Most DBA/2J insertions were also identified in DBA/1J (76/84). Similar to deletions, sensitivity (90.48%) of the SV insertion calling pipeline was similar to estimates reported previously in MGP strains [40,41].

Little is known about structural variation in the genomes of the 13 strains included in this study. However, many of the structural variants which we identified were also shared with another strain in the MGP variation catalogue. For example, a 6-kb deletion in *BFSP2* which results in alterations of optical lens quality has previously been identified in the FVB/NJ and 129 strains [43]. From our analysis, we identified this mutation in both the NZB/B1NJ and NZW/LacJ strains. Although neither of these strains are known models for examining optical lens quality, utilising either strain in studies which may rely on vision or lens quality could lead to spurious results.

Coding sequences for all genes annotated to the mouse genome were then compared against our deletion call set to identify genes which may have their function disrupted. A total of 4,954 structural deletions overlapped the coding sequence of 3,316 genes. Of these, 2,462 were novel and 2,110 were private to one of the 13 strains in our study. Similarly, 4,620 structural insertions overlapped the coding sequence of 5,076 genes, many of which were either novel (2,557) or private (2,366) to one of the 13 strains strains.

### RF/J strain enriched for private missense variants in cancer susceptibility genes

To investigate the potential biological mechanisms underlying disease phenotypes for which these strains are used to study, two analyses were carried out. The first study focused on the RF/J strain, which was developed at the Rockefeller Institute through breeding of the A, R and S strains [44]. Due to the natural predisposition of RF/J mice to develop myeloid leukemia, thymic lymphoma and reticulum cell sarcomas [44], this strain is an important model used to study cancer onset, progression and pathogenesis. This strain also has a propensity to develop lung, ovary and hematopoietic tumours. In particular, this strain is widely used to study the effects of radiation-induced myeloid leukemia due to its high induction rate (between 50-90%) [44].

To identify and prioritise candidate genes for further analysis, only genes which contained at least one missense variant private to the RF/J strain were considered for further analysis. A total of 122 genes were identified which met this condition (Additional file 7). Several of the genes in this list, such as *Adam15* [45], *Cdh1* [46], *Msh3* and *Kit* [47], are known candidates for roles in tumorigenesis and cancer pathogenesis. For example, mice nullizygous for the *Msh3* gene are reported to have a significantly higher spontaneous incidence rate of tumours than wild-type counterparts [48]. The *Msh3* gene is involved in the DNA repair mechanism, and plays an important role in mismatch recognition [49]. The mismatch repair pathway identifies and eliminates errors that arise during DNA replication, and defects in this mechanism can lead to the accumulation of mutations and cancer [50]. In fact, loss or silencing of the *Msh3* gene has been reported to frequently occur in several human cancers including lung [51] and breast [52]. In addition, associations of mutations in the *Msh3* gene to colon [53] and prostate [54] cancer in humans have been reported.

Another gene involved in the DNA damage response, *PALB2,* was also identified in our candidate set. Mutations of this gene are known to increase risk of several cancers including breast [55] and pancreatic cancer [56]. Mutations of *PALB2* are commonly identified in patients with Fanconi Anaemia. In fact, patients with *PALB2* related Fanconi Anaemia also have an increased predisposition to malignancies associated with acute myeloid leukemia [57] and Wilms tumours [58]. Interestingly, the *WT1* (Wilms tumour) gene was also identified in our candidate gene set as carrying a private missense SNP.

Additional genes of particular note in our candidate set included the tumour suppressor gene *Cdh1,* mutations in which are known to predispose to a wide range of cancer types [59], and the *Fas* gene. *Fas,* a member of the tumour necrosis factor receptor family of proteins, plays a key role in activation of apoptosis and has been implicated in the pathogenesis of several malignancies [60]. Dysregulation of apoptosis related pathways is also commonly observed in cancer, and defects of this signalling process are known to contribute to drug therapy insensitive tumours [61]. Furthermore, polymorphisms in the promoter of the *Fas* gene have been associated with an increased risk of developing acute myeloid leukemia in humans [62]. Interestingly, following Y-irradiation-induced thymic lymphoma, *Fas* expression has also been shown to be lower in the RF/J strain compared to the SPRET/EiJ and BALB/cJ strains both of which are known to be more resistant to this induced phenotype [63]. Although no private missense SNPs in this gene were identified in the BALB/cJ strain, one missense SNP in exon 2 was identified. Similarly, 17 missense SNPs were identified in the wild-derived SPRET/EiJ strain, 14 of which were private to this strain. Due to each of the above mentioned strains containing missense SNPs in this gene, it is difficult to determine the exact role this gene or mutations in this gene may play in the pathogenesis of cancer phenotypes observed in the RF/J strain.

To examine the overall effect of private missense mutations in RF/J, we identified biological pathways over-represented in this gene set. Significantly over-represented REACTOME [64] and KEGG [65] pathways were identified using the hypergeometric test available through the pathway over-representation analysis tools at InnateDB [66]. After correcting p-values using the benjamini-hochberg method, 4 pathways were significantly over-represented (p<05). Notably, this list included the KEGG pathway ‘Pathways in cancer’ (p=0.046). Each of the remaining 3 significantly over-represented pathways were related to extracellular matrix (ECM) functions (Table 3). In fact, the most significantly over-represented REACTOME and KEGG pathways were “Degradation of the extracellular matrix” (p=0.0049) and “ECM-receptor interaction” (p=0.013) respectively.

**Table 3.** **Significantly over-represented biological pathways and candidate genes identified in RF/J**. Candidate genes are those that are found in the respective pathways and contained a missense SNP private to the RF/J strain. Corrected p-values are based on the Benjamini-Hochberg method.

| Pathway Name | Database | p-value<br>(corrected) | Candidate genes |
| --- | --- | --- | --- |
| Degradation of the extracellular matrix | REACTOME | $4.91 \times 10^{-3}$ | *Adam15, Cdh1, Col4a4, Eln, Mmp11* |
| Extracellular matrix organization | REACTOME | $1.15 \times 10^{-2}$ | *Adam15, Cdh1, Col4a4, Eln, Lama3, Mmp11* |
| ECM-receptor interaction | KEGG | $1.32 \times 10^{-2}$ | *Col4a4, Itga10, Itga7, Lama3* |
| Pathways in cancer | KEGG | $4.56 \times 10^{-2}$ | *Cdh1, Col4a4, Fas, Kit, Lama3, Msh3* |

The ECM is a fundamental component of the cellular microenvironment, composed of a complex mix of structural and functional molecules [67]. The ECM has important roles in a diverse set of functions including cellular development, tissue differentiation, growth and tissue homeostasis [68]. In addition, components of the ECM are in continual contact with epithelial cells and mediate signalling of processes involved in regulation of cell proliferation, apoptosis and differentiation [69]. Notably, “ECM-receptor interaction” has previously been found to be enriched for genes differentially regulated in cancer cell lines compared to normal tissue [70], and in tumour expression patterns compared to normal tissue [71]. In addition, abnormal ECM has been implicated in the progression of cancer. In fact, altered ECM is commonly found in patients with cancer [72].

To investigate whether these results may be due to an annotation bias within KEGG or REACTOME, the same pathway over-representation analysis was performed for each of the 13 strains' private missense mutations (Additional file 8). No strain's gene set was enriched for genes involved in “Pathways in cancer”, “Degradation of the extracellular matrix” or “Extracellular matrix organization”. However, “ECM-receptor interaction” was also significantly over-represented in the MOLF/EiJ gene set (p=0.0026). Though it may not be appropriate to carry out such analysis using wild-derived strains such as MOLF/EiJ due to the relatively low representation of wild-derived strains in the MPG catalogue compared to the classical inbred strains. For example, approximately 56 times as many private SNPs were identified in MOLF/EiJ compared to RF/J (Table 1). Consequently, there is nearly 28 times as many genes which met the inclusion criteria using MOLF/EiJ (3,366 genes) when compared to the RF/J gene set (122 genes).

### Collagen-induced arthritis candidate genes and biological pathways

The second study carried out focused on the genetic and phenotypic differences between the genetically similar DBA/1J and DBA/2J strains. The DBA strain was developed by Clarence C Little in 1909 and is the oldest of all inbred mouse strains. During 1929 and 1930, crosses between substrains lead to the establishment of several new substrains including DBA/1J and DBA/2J [34]. Just over 90.72% of SNP sites identified in the DBA/1J strain are identical to the DBA/2J strain, however, these strains exhibit several phenotypic differences including total body weight and body fat weight [73]. In addition, differences in circulating plasma levels of triglycerides, high-density lipoprotein, leptin and insulin have been reported between DBA/1J and DBA/2J [34]. Notably, these strains also display divergent susceptibility to collagen induced arthritis (CIA), a model for human rheumatoid arthritis. The DBA/1J strain, sequenced as part of this study, is susceptible to CIA whereas the DBA/2J strain is resistant [74]. To identify potential variation, and genes that may be involved in these divergent phenotypes, all SNPs identified in DBA/1J which are shared with the DBA/2J strain were removed. This meant we were left with a set of DBA/1J specific SNPs. However, despite the genetic similarity of these strains, they have different haplotypes at the MHC locus [75]. To remove SNPs in this region not associated with CIA susceptibility, all SNPs shared with the CIA resistant but MHC identical FVB/NJ strain were also removed. A total of 258,416 SNPs remained in this DBA/1J specific CIA susceptibility candidate set. From this, all genes containing at least a single missense variant with the additional SIFT annotation of “deleterious” were identified (123 SNPs identified in 101 genes). Similar to the previous study, this gene set was then used to identify significantly over-represented biological pathways using the tools available from InnateDB [66].

Murine collagen induced arthritis, a widely used model for human rheumatoid arthritis (RA), is characterised as an autoimmune inflammatory joint disease which primarily affects synovial joints ultimately leading to joint impairment and disability [14]. The pathogenesis of RA involves the influx of both innate and adaptive immune cells, which in turn promote pro-inflammatory cytokine production and decreased synthesis of anti-inflammatory cytokines [76]. Normally the inflammatory response is balanced by the release of anti-inflammatory molecules which prevent chronic inflammation. Disruption of the anti-inflammatory balancing mechanism can lead to sustained chronic inflammation as observed in rheumatoid arthritis. Interestingly, our list of candidate genes containing a missense deleterious SNP also included the anti-inflammatory mediator interleukin–10 receptor a subunit *(IL10RA).* Interleukin–10 *(IL10)* is an antiinflammatory cytokine whose functions are mediated by binding to *IL10RA* which in turn stimulates signalling cascades used to inhibit production of pro-inflammatory cytokines such as tumour necrosis factor a *(TNFA)* [77]. *IL–10*DBA/1J mice have increased CIA severity compared to mice either homozygous or heterozygous at the same locus [78]. Similarly, therapeutic treatment with *IL-10* has been shown to suppress the development of CIA in rats [79]. In addition, mutations in the *IL10RA* have been implicated in the pathogenesis of several inflammatory autoimmune diseases such as murine lupus [80] and ulcerative colitis [81]. Diminished *IL10* signalling, through mutations in the *IL10RA,* have been implicated in the severity of several autoimmune diseases including inflammatory bowel disease and arthritis [81,82].

Similar to RF/J, pathway over-representation analysis was carried out using the candidate gene set identified in the DBA/1J analysis. A total of 10 pathways were significantly over-represented (corrected p<0.05), many of which are related to innate immunity and inflammation signalling (Table 4). The most significantly over-represented pathway was “cellular adhesion molecules (CAMs)” (p=3.98 × 10^−4^). Cell adhesion molecules are expressed on the surface of cells and play a critical role in the inflammatory response. This pathway has been shown to be enriched for genes with a differentially methylated signature in synovial tissue of RA patients compared to patients without RA [83]. Several additional KEGG pathways identified in our analysis have also been implicated in the pathogenesis of RA including “cytokine-cytokine receptor interaction” (p=0.047) and “antigen processing and presentation” (p=0.011) [84]. Notably, none of the candidate genes identified in the cytokine-cytokine receptor interaction pathway are shared with candidate genes identified in either CAMs or antigen processing and presentation pathways. Whereas all candidates identified in the antigen processing and presentation pathway are also found in the CAMs pathway. Given the complex nature of RA, it is not surprising that many genes and biological pathways contribute to the pathogenesis of RA [85]. Complimenting evidence from individual candidate gene lists with biological information available through interaction and pathway databases may help identify new targets for therapeutic treatments of complex diseases such as RA. In spite of this, additional work utilising other high-throughput technologies (e.g. RNA-seq and proteomics) and *in vivo* models is still required to fully understand the onset and severity of CIA and ultimately RA.

**Table 4.** **Significantly over-represented biological pathways and candidate genes identified in DBA/1J.** Candidate genes are genes that are found in the overrepresented pathways and which contained a missense, deleterious SNP found in DBA/1J and not in DBA/2J or FVB/NJ. Corrected p-values were calculated using the Benjamini-Hochberg method.

| Pathway Name | Database | p-value<br>(corrected) | Candidate genes |
| --- | --- | --- | --- |
| Cell adhesion molecules (CAMs) | KEGG | $3.98 \times 10^{-4}$ | *Esam, H2-M10, H2-M10, H2-T22, Itga9, Sdc3* |
| Type I diabetes mellitus | KEGG | $7.54 \times 10^{-3}$ | *H2-M10, H2-M10, H2-T22* |
| Graft-versus-host disease | KEGG | $7.59 \times 10^{-3}$ | *H2-M10, H2-M10, H2-T22* |
| Autoimmune thyroid disease | KEGG | $8.55 \times 10^{-3}$ | *H2-M10, H2-M10, H2-T22* |
| Allograft rejection | KEGG | $1.01 \times 10^{-2}$ | *H2-M10, H2-M10, H2-T22* |
| Antigen processing and presentation | KEGG | $1.06 \times 10^{-2}$ | *H2-M10, H2-M10, H2-T22* |
| Viral myocarditis | KEGG | $1.14 \times 10^{-2}$ | *H2-M10, H2-M10, H2-T22* |
| Endocytosis | KEGG | $3.07 \times 10^{-2}$ | *Arap3, H2-M10, H2-M10, H2-T22* |
| Linoleic acid metabolism | KEGG | $4.49 \times 10^{-2}$ | *Cyp2c44, Cyp2c50* |
| Cytokine-cytokine receptor interaction | KEGG | $4.73 \times 10^{-2}$ | *Il10ra, Il28ra, Tnfrsf1b, Tnfrsf8* |

## Conclusions

We report the significant expansion of the Mouse Genomes Project resource to incorporate deep whole-genome sequencing of 36 inbred laboratory strains. The amount of SNP variation found has increased from 56M to 71M sites, comparable in genetic variation to the final 1000 Genomes set [86]. In this paper, we focus on the 13 new laboratory strains that include further representatives of the classical laboratory and wild-derived strains. The original project sought to sequence as widely possible from the laboratory mouse subspecies groups. These 13 new strains mean that we now have several examples of highly related pairs of strains that are known to display divergent phenotypes, which could accelerate the identification of causative variants.

We focus on two specific examples of how the variation data detected can be used to prioritise genes of interest for well known phenotypes in the new strains. In the first example, we focus on the RF/J strain which has a highly penetrant predisposition to develop acute myeloid leukemia, thymic lymphoma and reticulum cell sarcomas. We used the expanded variation catalog to identify candidate genes containing missense mutations that are private to the RF/J strain. Several notable genes with a known role in cancer were identified in our candidate gene set (e.g. *Msh3, Kit, Fas).* This gene set was then further interrogated leading to the identification of biological pathways with potential roles in cancer susceptibility in this strain. In the second analysis, we investigate the genetic basis of collagen induced arthritis (a model of human rheumatoid arthritis) using two genetically very similar strains with divergent phenotypes of susceptible (DBA/1J) and resistant (DBA/2J). Through comparison of variants identified in both strains, we were able to identify a candidate gene set of 101 genes involved in the susceptibility of DBA/1J to this induced phenotype. Further analysis of this gene set revealed several biological processes involved in immune signalling and inflammation which may play a role in the pathogenesis of human rheumatoid arthritis. When using the mutation catalogs for genetic mapping studies, it must be noted that the structural variation calls have an error rate of up to 10%. A recent study found that more accurate structural variation calls can be derived from long read technologies compared to those produced by short read detection methods [87], therefore we expect that future releases of variants on these strains will be even more accurate.

These strains represent a significant increase in the disease models available in the MGP variation catalogue which will enable improved dissection of the genetic variation underlying many human diseases. The expansion of the catalog of sequenced strains means that we can identify mutations that are more likely to be truly private to individual strains thereby reducing the size of candidate variant sets. Sequencing of genetically diverse strains, such as the wild-derived MOLF/EiJ, are invaluable resources for identifying new genetic variation not present in classical inbred laboratory strains. However, in wild-derived strains many loci are composed of novel alleles and divergent haplotypes from the current reference genome [88]. Efforts are currently ongoing to create *de novo* assembled genomes for several classical and wild-derived strains [89] which will enable more accurate examination of the genomic structure and genetic variation responsible for phenotypes observed in these strains.

## Methods

### DNA sequencing and read alignment

DNA from a single male of each inbred mouse strain was obtained from the Jackson Laboratories. Jackson Laboratory stock numbers for each strain are contained in Table s1 of Additional file 1. DNA was sheared to 200-300bp fragment size using a Covaris S2. A single illumina TruSeq v3 sequencing library was created for each strain according to manufacturer's protocols. Each library was sequenced on a HiSeq 2000 over four lanes. Each lane was genotype checked using Samtools/Bcftools v1.2 (http://www.htslib.org) ‘gtcheck’ command by comparing variant sites against the mouse Hapmap [22]. Library and sequencing statistics for each sample are available in Table s3 of Additional file 1.

Sequencing reads from each lane were aligned to the C57BL/6J GRCm38 (mm10) mouse reference genome [5] using BWA-MEM (v0.7.5) [90] with default parameters. For each library, aligned reads from each lane were merged using Picard Tools (v1.64) [91], resulting in a single BAM file corresponding to all aligned reads for a single strain. Each BAM file (one for each strain) was then sorted and filtered for possible PCR and optical duplicates using Picard Tools (v1.64). To improve SNP and indel calling, the GATK v3.0 “IndelRealigner” tool [92], was used to realign reads around indels using default options.

### SNP and indel discovery

SNPs and indels were identified using a combination of SAMtools mpileup (v1.1) [93] and BCFtools call (v1.1). SAMtools mpileup captures summary information from input BAMs, and uses alignment information to estimate genotype likelihoods. BCFtools, based on the VCFtools package [94], then uses this information, with the inclusion of a prior, to perform the variant calling. The following options were specified for SAMtools; ‘-t DP,DV,DP4,SP,DPR,INFO/DPR-E-Q 0-pm3-F0.25-d500’ and BCFtools call; ‘-mv-f GQ,GP-p 0.99’. To improve accuracy of indel calls, indels were then left-aligned and normalised using the bcftools norm function with the parameters ‘D-s-m+indels’.

To ensure high quality variants were only used for later analysis, bcftools annotate was used to soft filter SNPs and indels identified. These filters are used to identify low quality variant calls, and remove false SNP and indel calls due to alignment artifacts. Only high quality (i.e. passed all filters) homozygote variants were retained. Filter values and cut-offs used to identify low quality variants are described in Table s4 of Additional file 1.

### SNP and indel annotation and prediction of consequences

Using mouse gene models obtained from Ensembl 78, the Ensembl Variant Effect Predictor tool (version 78) [23] was used to predict the effect of SNPs and indels on gene transcripts. VEP assigns variants to functionally relevant categories based on the predicted effect that a variant has on a transcript or gene. Consequences assigned to variants within coding regions of a gene include stop losses/gains, missense variants and splice variants. Notably, a single variant may have different estimated consequences between alternative transcripts of the same gene. To predict whether amino acid substitutions would be damaging, Grantham matrix scores (GMS) [29] were calculated for each variant. Grantham matrix scores are used to predict the effect of substitutions between amino acids. Variants with a calculated GMS between 100 and 150 (100 < GMS ≤ 150) were considered moderately radical, and variants with a GMS > 150 were considered radical changes. Additionally, SIFT [95] scores were estimated for all variants. SIFT predicts amino acid substitution effects on protein function, and classifies variants as being either tolerated or deleterious.

### Structural variation calling

A combination of BreakDancer [96], cnD [97] and LUMPY [98] were used to identify larger structural deletions/losses (greater than 100bp). Breakdancer identifies large deletions using deviations from the expected mate pair distance and cnD identifies losses of genomic segments using changes in mapping depth across regions. LUMPY utilises multiple SV signals, including read-pair and split-read information to identify potential SV breakpoints. All parameters used by breakdancer, cnD and LUMPY are described in Table s5 of Additional file 1. The deletion call sets for each strain were then collapsed into strain-specific non-redundant sets of deletions by merging deletions with a reciprocal overlap greater than 90%. This meant that for each merged deletion there were two possible deletion breakpoints, the primary breakpoints, which are those identified by breakdancer, cnD or LUMPY, and a secondary breakpoint, which are the outer bounds of merged deletions.

To identify SV insertions, RetroSeq (transposable element insertions) [99], cnD (copy number gains) [97], scalpel [100] and manta (both novel sequence insertions) [101] were used. Parameters for each analysis are contained in Table s5 of Additional file 1.

Calls made by each tool were then filtered to remove any SV within 500bp of an assembly gap, or 20kb of a telomere or centromere as mapping artifacts in these regions can cause false structural variant calls. Additionally, SVs greater than 1MB in size were removed as these were generally false positive calls due to poor mapping among reads.

### Identification and count of private variation

Variants identified in the 13 strains examined in this study were then incorporated in the mouse genomes variation catalogue. The MGP variation catalogue now contains variants identified in 36 different inbred laboratory mouse strains. To obtain a list of high confidence variants private to each strain, sequencing data from the remaining 23 strains contained in the MGP variation catalogue was aligned to the mouse reference genome using the same pipeline as described above.

Similarly, the same SAMtools and BCFtools pipeline as above was used to identify SNPs and short indels. SNP and indel locations from all 36 strains were collated into a single file which was used to generate genotypes at those locations in all strains using samtools mpileup with the −l option enabled (i.e. report the genotype at all locations listed). The same bcftools call parameters as the initial variant calling step were used. This allowed the identification of high quality homozygous reference locations, as well as low quality regions and regions with no sequence information
(uncallable regions).

Private SNPs were identified as locations in which a single high quality homozygote alternative variant was called in only one of the 36 strains. In the remaining 35 strains, the location must be a high quality homozygous reference genotype allowing a maximum of 5 low quality or missing genotypes in any of the strains. Similarly, private indels were identified for each strain. A private indel was defined as any high quality homozygote alternative indel unique to a single strain.

To identify SV deletions that may be the result of the same ancestral event, deletions were first identified in each of the remaining 23 MGP strains using the same approach as described above. Deletions identified in each individual strain were compared with the deletions identified in all other strains (13 strains sequenced as part of this study and the additional 23 MGP strains). A non-redundant across-strain deletion set was identified, as before, using a greater than 90% reciprocal overlap to merge deletions. Following this merge, private SV deletions were identified as those that did not overlap with another deletion, and thus were not merged with any other deletion.

Similarly, as with predicted SV deletions, SV insertions were identified for all 36 strains contained in the MGP variation catalogue, including the 13 strains from this study. The insertion call sets for each strain were then merged, and used to identify insertions private to each strain.

### Variants calls accuracy

To assess the accuracy of SNP calls made for each strain, genotypes were obtained from the mouse HapMap resource [22] for each of our strains and compared with the corresponding SNP identified in our catalogue for each of the 13 strains separately. The mouse HapMap contains approximately 138,000 homozygous sites genotyped in 94 inbred laboratory mouse strains. Genotype locations for each of the 13 strains were mapped from mm9 to mm10 of the reference genome, and any missing genotypes were removed from further analysis. Between 115,572 (MOLF/EiJ) and 129,825 (C58/J) remained after mapping SNP locations from mm9 to mm10 and removing missing genotypes. Sensitivity of the SNP calling pipeline was estimated by calculating the number of genotypes concordant between the HapMap and the variation catalogue for each of the 13 strains. The number of genotypes from the HapMap available for comparison with each of the 13 strains and the respective concordance rate are contained in Additional file 4.

A dataset of PCR validated SV insertions and deletions for 7 MGP strains (AKR/J, A/J, BALB/cJ, DBA/2J, CBA/J, C3H/HeJ and LP/J) were obtained from Yalcin *et. al.* [39]. Coordinates for all variants were first mapped from NCBIm37 to GRCm38. Using the SV calling pipeline described above, sequencing data for these 7 strains was used to call structural variants that could be compared to the PCR validated set to assess the accuracy of our SV calling pipeline. In addition, SVs identified in the DBA/1J from our study were compared to the PCR validated variants for DBA/2J (Table s2 of Additional file 1).

### Identification of candidate genes through prioritisation of damaging SNPs

To demonstrate the usefulness of this data, two studies investigating the genetic basis of cancer susceptibility and collagen induced arthritis (CIA) were carried out. The first investigation involved identifying all genes that contained a missense variant that was private to the RF/J strain. This strain is an important model used to investigate the onset and progression of several cancers including leukemia and reticulum cell sarcomas. This gene set was then used to identify any biological pathways that may be enriched for genes from this set (described below).

The second investigation was carried out to investigate the genetic basis of CIA. Two strains, DBA/1J and DBA/2J, were chosen as they are genetically similar (>90.72% of DBA/1J SNPs are shared with DBA/2J) but phenotypically divergent for susceptibility to collagen induced arthritis; DBA/1J is susceptible however DBA/2J is resistant. All SNPs identified in these two strains were collated into a single file. The SNPs specific to DBA/2J or shared with DBA/2J were then removed leaving a DBA/1J specific set of SNPs. However, these strains differ at the MHC locus. To account for the differing MHC alleles, SNPs shared with the CIA resistant FVB/NJ strain were also removed. This SNP set was then filtered to only include missense SNPs with a SIFT annotation of deleterious. All genes containing these variants were then identified, and this gene set was used to identify biological pathways that may be involved in CIA susceptibility.

### Pathway over-representation analysis

For each of the candidate gene sets described above, a pathway overrepresentation analysis was performed to identify biological pathways or processes that are statistically enriched for this gene set. This was done using the pathway overrepresentation tools available from InnateDB [66], and analysed pathway annotations available from the REACTOME [64] and KEGG [65] databases. This test was based on the hypergeometric algorithm, and correction of p-values was done using the Benjamini-Hochberg method. Significantly over-represented pathways were defined as those with a corrected p<0.05.

## Abbreviations

AML; Acute Myeloid Leukemia. bp; base pair. CDS; Coding Sequence. CIA; Collagen Induced Arthritis. GATK; Genome Analysis ToolKit. GMS; Grantham Matrix Scores. Indel; insertion/deletion polymorphism. MGP; Mouse Genomes Project. PCR; Polymerase Chain Reaction. SNP; Single Nucleotide Polymorphism. SV; Structural Variant. UTR; Untranslated Region.

## Declarations

### Availability of data and material

All raw sequencing reads are available from the European Nucleotide Archive (ERP000927; see Table s1 in Additional file 1 for sample identifiers). The sequence variants are available from the European Variation Archive (PRJEB11471) and the Mouse Genomes Project ftp site (ftp://ftp-mouse.sanger.ac.uk/).

### Author contributions

KWH, JF, DJA, and TMK initiated the ideas for the study. AJD, KW and TMK carried out the computational analysis and drafted the manuscript. All authors read and approved the manuscript.

### Ethics approval and consent to participate

DNA provided by The Jackson Laboratories was obtained in accordance with local ethical guidelines and regulations.

## Acknowledgements

Not applicable.

## Additional files

### Additional file 1

**Format**: PDF.

**Title**: Supplemental tables.

**Description**: Supplemental tables s1-s5.

### Additional file 2

**Format**: PDF.

**Title**: SNP, indel and SV density plots for all strains.

**Description**: SNP, indel and SV (deletions and insertions) density plots for all chromosomes (1-19, X and Y) for each strain.

### Additional file 3

**Format**: XLSX.

**Title**: Pairwise variant comparisons.

**Description**: The number of SNPs and indels shared between any two strains contained in the MGP variation catalogue.

### Additional file 4

**Format**: CSV.

**Title**: Per strain genotype concordance with HapMap.

**Description**: Total HapMap positions available for comparison with each strain, and the corresponding number of concordant sites.

### Additional file 5

**Format**: XLS.

**Title**: VEP, GMS and SIFT annotations for SNPs and indels in each of the 13 strains.

**Description**: Total number of SNPs and indels annotated into each functional consequence predicted by VEP. Total number of SNPs with an estimated moderate (using GMS), radical (GMS), tolerated (using SIFT) and deleterious (SIFT) effect is also provided.

### Additional file 6

**Format**: .txt.gz (gzip compressed text file).

**Title**: Private SNPs with multiple VEP consequences.

**Description**: Private SNPs with multiple predicted VEP consequences for each strain strain. Consequence predictions are based on gene models obtained from Ensembl 78.

### Additional file 7

**Format**: CSV.

**Title**: Candidate genes identified in the RF/J analysis.

**Description**: All genes which contained at least one missense SNP only found in the RF/J strain.

### Additional file 8

**Format**: CSV.

**Title**: Per strain over-represented pathways using genes containing private missense SNPs.

**Description**: For each strain separately significantly over-represented pathways were identified using only the genes which contained at least one private missense variant.

